# The mRNA cap 2’O methyltransferase CMTR1 regulates the expression of certain interferon-stimulated genes

**DOI:** 10.1101/2020.03.05.980045

**Authors:** Graham D. Williams, Nandan S. Gokhale, Daltry L. Snider, Stacy M. Horner

**Author notes:** To whom correspondence should be addressed: Stacy M. Horner, Ph.D., Department of Molecular Genetics and Microbiology, Duke University Medical Center, 213 Research Dr., Box 3053 DUMC, Durham, NC USA 27710.

## Abstract

Type I interferons (IFN) initiate an antiviral state through a signal transduction cascade that leads to the induction of hundreds of IFN-stimulated genes (ISGs) to restrict viral infection. Recently, RNA modifications on both host and viral RNAs have been described as regulators of infection. However, the impact of host mRNA cap modifications on the IFN response and how this regulates viral infection is unknown. Here, we reveal that CMTR1, an ISG that catalyzes 2’O methylation of the first transcribed nucleotide in cellular mRNA (Cap 1), promotes the protein expression of specific ISGs that contribute to the antiviral response. Depletion of CMTR1 reduces the IFN-induced protein levels of ISG15, MX1, and IFITM1, without affecting their transcript abundance. However, CMTR1 depletion does not significantly affect the IFN-induced protein or transcript abundance of IFIT1 and IFIT3. Importantly, knockdown of IFIT1, which acts with IFIT3 to inhibit the translation of RNAs lacking Cap 1 2’O methylation, restores protein expression of ISG15, MX1, and IFITM1 in cells depleted of CMTR1. Finally, we found that CMTR1 plays a role in restricting RNA virus replication, likely by ensuring the expression of specific antiviral ISGs. Taken together, these data reveal that CMTR1 is required to establish an antiviral state by ensuring the protein expression of a subset of ISGs during the type I IFN response.

**Importance:** Induction of an efficient type I IFN response is important to control viral infection. We show that the host 2’O methyltransferase CMTR1 facilitates the protein expression of ISGs in human cells by preventing IFIT1 from inhibiting the translation of these mRNAs lacking cap 2’O methylation. Thus, CMTR1 promotes the IFN-mediated antiviral response.

## Introduction

Interferons (IFN) are cytokines that establish an antiviral cellular state through transcriptional induction of interferon-stimulated genes (ISGs) (1). Following engagement of IFN (type I and type III) with its cognate cell surface receptors, a JAK-STAT signaling cascade is activated that results in the formation of the ISGF3 transcription factor complex that binds to IFN response elements in the promoters of ISGs. Following their transcription, ISG mRNAs are translated, generating the proteins that restrict viral infection and viral spread (2, 3). Full induction of this antiviral state requires hundreds of ISGs to be upregulated efficiently, and therefore cells must encode mechanisms to ensure their regulated expression. Such regulation can occur at both the transcriptional and post-transcriptional level (4-6). While several post-transcriptional controls of antiviral cytokines and signaling molecules have been defined, similar mechanisms that regulate ISG expression have been largely unexplored (5, 7).

Chemical modifications at the 5’ end of mRNAs are required for optimal gene expression (8). One such modification is N^7^-methylguanosine (m^7^G) that is added co-transcriptionally to the first transcribed nucleotide at the 5’ end of mRNAs (referred to as Cap 0; **Figure 1A**). In higher eukaryotes, the ribose 2’O hydroxyl group of the first transcribed nucleotide is methylated, resulting in the Cap 1 structure (**Figure 1A**) (8). Approximately 50% of cellular mRNAs are also methylated at the ribose 2’O hydroxyl group of the second transcribed nucleotide (9). These mRNA cap modifications have been shown to have a wide variety of functions. For example, the m^7^G modification in the mRNA cap recruits the translation initiation machinery for cap-dependent translation, and Cap 1 2’O methylation can further enhance this translation (8, 10, 11). mRNA cap 2’O methylation has also been shown to prevent degradation of modified transcripts by exoribonucleases or serve as a motif that marks mRNA as “self” to prevent recognition by the IFN-induced RNA binding proteins RIG-I and IFIT1 (12-15). RIG-I binding to Cap 0 mRNA can activate a signaling cascade that induces an IFN response (12, 16), whereas IFIT1 binding to Cap 0 mRNA inhibits its translation (13, 17-19). This translational inhibition by IFIT1 is in part mediated by its interaction with the IFN-induced protein IFIT3, which increases the specificity of IFIT1 for Cap 0 mRNAs (20-22). As such, many viruses have evolved mechanisms to evade RIG-I and IFIT1 sensing of their 5’ RNA ends. Some examples of these viral strategies to evade RIG-I and IFIT1 sensing include encoding proteins that bind to and shield Cap 0 in viral mRNA; acquiring Cap 1 from cellular mRNAs; altering their RNA structures at the 5’ end; or encoding their own viral Cap 1 2’O methyltransferases (8, 23-30), highlighting the overall importance of this RNA modification in “self vs non-self” recognition.

**Figure 1:**
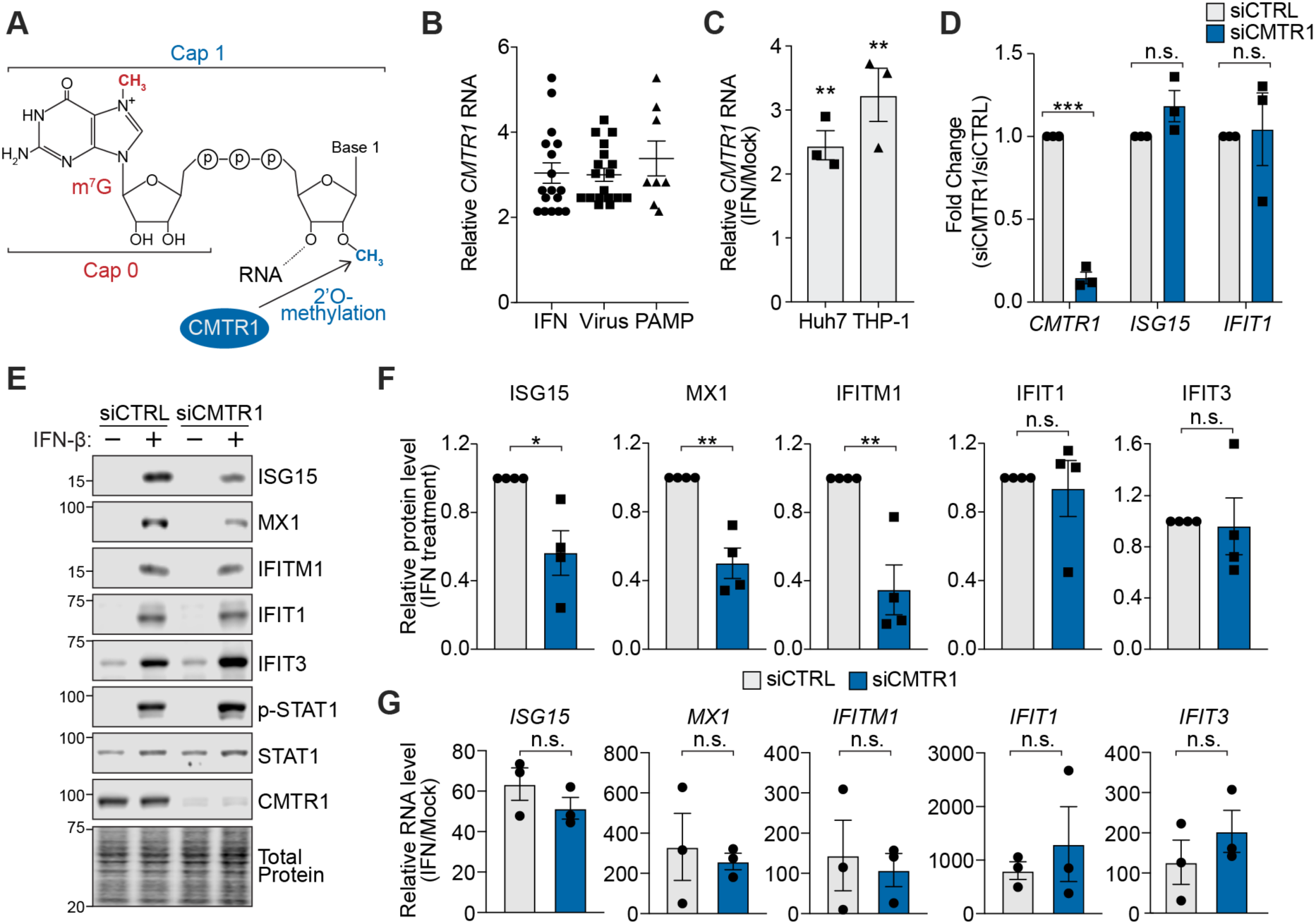
CMTR1 promotes the protein expression of specific ISGs. **(A)** Schematic of 5’ cap structure, with m^7^G (Cap 0) and 2’O methylation of the first transcribed nucleotide catalyzed by CMTR1 (Cap 1) **(B)** Fold change of *CMTR1* treated with the indicated stimuli (type I, type II, or type III IFNs; viruses (human rhinovirus 16, Newcastle disease virus, human cytomegalovirus strain TB40E, influenza A virus, yellow fever virus 17D); pathogen-associated molecular patterns (PAMP: lipopolysaccharide or poly (I:C)) relative to untreated samples. Data were obtained from EMBL-EBI Expression Atlas RNA-seq database (34). **(C)** RT-qPCR analysis of *CMTR1* relative to *GAPDH* in either Huh7 or THP-1 cells following IFN-β treatment, as compared to levels in untreated cells. **(D)** RT-qPCR analysis of *CMTR1, ISG15*, and *IFIT1* relative to *GAPDH* in mock-treated Huh7 cells transfected with the indicated siRNAs. **(E)** Representative immunoblot of extracts from mock- or IFN-β-treated Huh7 cells transfected with the indicated siRNAs. **(F)** Quantification of the immunoblots from (E), normalized to total protein and graphed relative to siCTRL. **(G)** RT-qPCR analysis of *ISGs* relative to *GAPDH* in IFN-β-treated Huh7 cells transfected with the indicated siRNAs. Data are graphed as the fold change relative to mock-treated cells. All IFN-β treatments were performed at 50 U/mL for 6 h. Values are the mean ± SEM of 3 (C, D, G) or 4 (F) biological replicates. * p < 0.05, ** p < 0.01, *** p < 0.001 by unpaired Student’s t test. n.s. = not significant.

Cap 1 2’O methylation of host transcripts is catalyzed by the cellular enzyme CMTR1 (31, 32). Interestingly, CTMR1, also called ISG95, is transcriptionally induced following induction of IFN (32, 33). Therefore, we hypothesized that CMTR1 may be required to promote the IFN-mediated induction of ISGs and the antiviral response. Here, we demonstrate that CMTR1 is required for full expression of specific ISGs in response to type I IFN. We found that depletion of CMTR1 results in decreased protein production of certain ISGs without affecting the abundance or stability of their associated transcripts. Importantly, this reduced ISG expression in the absence of CMTR1 leads to increased virus replication. In addition, we found that the inhibition of protein production of the CMTR1-regulated ISGs is mediated by the 2’O methylation sensor IFIT1, as loss of IFIT1 rescued their protein expression following CMTR1 depletion. Together, these results reveal that CMTR1 is required for efficient expression of ISGs and the resulting antiviral state.

## Results

### CMTR1 promotes the protein expression of specific ISGs

Previous studies have shown that CMTR1 is an ISG that is upregulated in response to multiple stimuli, including type I and type II IFN, viral infection, and multiple pathogen-associated molecular patterns (PAMPs) (**Figure 1B**) (32-34). We confirmed that the *CMTR1* transcript is induced in response to IFN-β in human Huh7 liver hepatoma cells and human monocyte THP-1 cells, as measured by RT-qPCR (**Figure 1C**). Because of the previously established role of CMTR1 in regulating gene expression, we hypothesized that it may regulate the expression of other genes induced by IFN. To test this, we depleted CMTR1 using siRNAs, and then measured the transcript levels and protein expression of a set of ISGs in the presence or absence of IFN-β. While others have shown that CMTR1 depletion induced *IFIT1* in the absence of exogenous IFN-β in primary human fibroblasts, we found that in unstimulated Huh7 cells, CMTR1 depletion did not induce ISG transcripts (**Figure 1D**) (16). However, we did find that CMTR1 depletion resulted in decreased protein expression of the ISGs ISG15, MX1, and IFITM1, while the expression of the ISGs IFIT1 and IFIT3 remained largely unaffected (**Figure 1E-F**). We also found that in Huh7 cells the relative RNA levels of these ISGs in response to IFN-β were not altered by CMTR1 depletion, as measured by RT-qPCR **(Figure 1G)**. These data reveal that CMTR1 promotes the protein, but not mRNA expression, of certain ISGs in response to IFN.

### CMTR1 is required for the antiviral response

Infection of human cells by RNA viruses is often restricted by the antiviral functions of the proteins encoded by ISGs (2, 3). Because CMTR1 is required for the expression of specific ISGs, we hypothesized that it would also regulate the antiviral response to IFN-sensitive viruses, such as the positive-sense RNA viruses Zika virus (ZIKV) and dengue virus (DENV) (35). Importantly, since both of these viruses encode their own Cap 1 2’O methyltransferase activity, they are not reliant on CMTR1 for 2’O methylation of their mRNA (36). At 48 hours post-infection with either ZIKV or DENV, we measured virus replication by using three complementary assays: the production of infectious virus in the supernatant by focus forming assay, viral RNA replication by RT-qPCR, and the percentage of infected cells by immunofluorescence microscopy. We found for both ZIKV and DENV, depletion of CMTR1 resulted in higher levels of infectious virus, higher levels of viral RNA, and a higher percentage of infected cells, as compared to control cells (**Figure 2A-C**). Further, depletion of CMTR1 resulted in increased viral spread as compared to control cells, in which viral replication was limited to discrete foci (**Figure 2C**). To determine directly whether CMTR1 contributed to viral restriction by type I IFN, we measured the ability of type I IFN to restrict infection by another IFN-sensitive virus, vesicular stomatitis virus (VSV), a negative-sense RNA virus which also encodes its own Cap 1 2’O methyltransferase (37-39), following CMTR1 depletion. In IFN-β treated Huh7 cells, CMTR1 depletion resulted in an increase in the percentage of VSV-infected cells as compared to control cells. Importantly, this increase was not seen in the absence of STAT1, which is required for IFN-mediated induction of ISGs (**Figure 2D)**. Together these results suggest that CMTR1 is required to facilitate the IFN-mediated induction of the cellular antiviral state.

**Figure 2:**
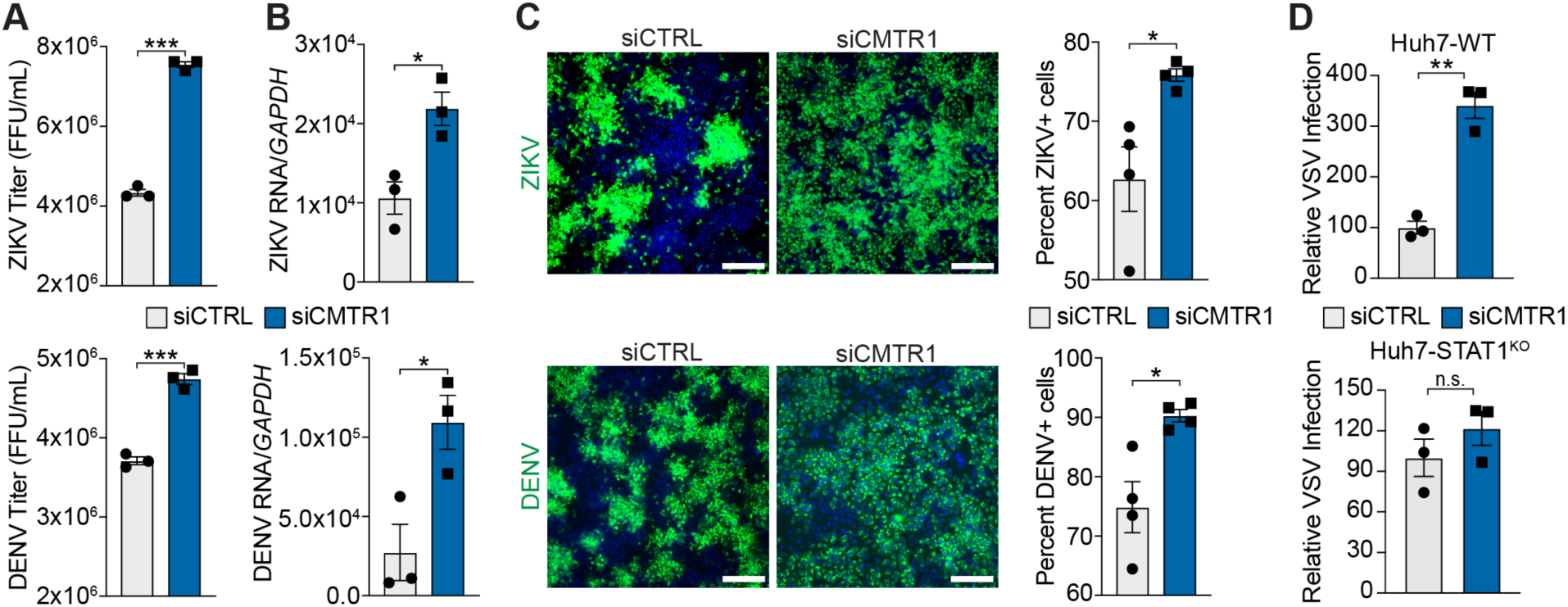
CMTR1 is required for the antiviral response. **(A)** Focus-forming assay of supernatants or **(B)** RT-qPCR analysis of viral RNA, relative to *GAPDH*, harvested from ZIKV- or DENV-infected Huh7 cells (48 hours post-infection (hpi); multiplicity of infection (MOI) 0.01) treated with the indicated siRNAs. **(C)** (Left) Representative fields of ZIKV- and DENV-infected Huh7 cells (48 hpi; MOI 0.01) treated with the indicated siRNAs and immunostained with anti-flaviviral E protein (green). Nuclei were stained with DAPI (blue). Scale bar: 100 μm. (Right) Quantification of the percentage of ZIKV- and DENV-infected Huh7 cells. **(D)** Quantification of the relative number of VSV-GFP-positive Huh7 WT and STAT1^KO^ cells after siRNA treatment and IFN-β pre-treatment (25 U/mL; 16 h) at 8 hpi and set to 100. For (C) and (D): ≥5000 cells were counted in each experiment per condition. Values are the mean ± SEM of 3 biological replicates. * p < 0.05, ** p < 0.01, *** p < 0.001 by unpaired Student’s t test. n.s. = not significant.

### CMTR1 does not regulate the nuclear export, RNA stability, or polysome association of specific ISGs

As we found that CMTR1 depletion results in lowered protein, but not mRNA expression of specific ISGs (ISG15, MX1, and IFITM1), we sought to determine the molecular mechanism underlying this change in protein expression. Cap 1 has been shown to shield mRNAs from degradation, regulate mRNA nuclear export, and promote mRNA translation (8, 9, 14, 15, 40). Therefore, we tested how loss of CMTR1 regulated these processes for ISG15, MX1, and IFITM1. Following IFN-β treatment, neither the nuclear export of these ISGs nor their stability were altered by CMTR1 depletion (**Figure 3A-B**). To determine whether CMTR1 depletion affected mRNA translation, we performed polysome analysis of IFN-β-treated Huh7 cells. We found no difference in the overall polysome profiles of these cells following CMTR1 depletion, as compared to control cells (**Figure 3C**). Surprisingly, we also found no difference in the polysome association of the CMTR1-regulated ISG transcripts (*ISG15, MX1*, and *IFITM1*) following CMTR1 depletion (**Figure 3D)**. Collectively, these results suggest that loss of CMTR1 does not globally alter cellular translation in response to IFN-β, similar to results described by others (41), nor does it alter the nuclear export, transcript stability, or polysome association of the CMTR1-regulated ISGs.

**Figure 3:**
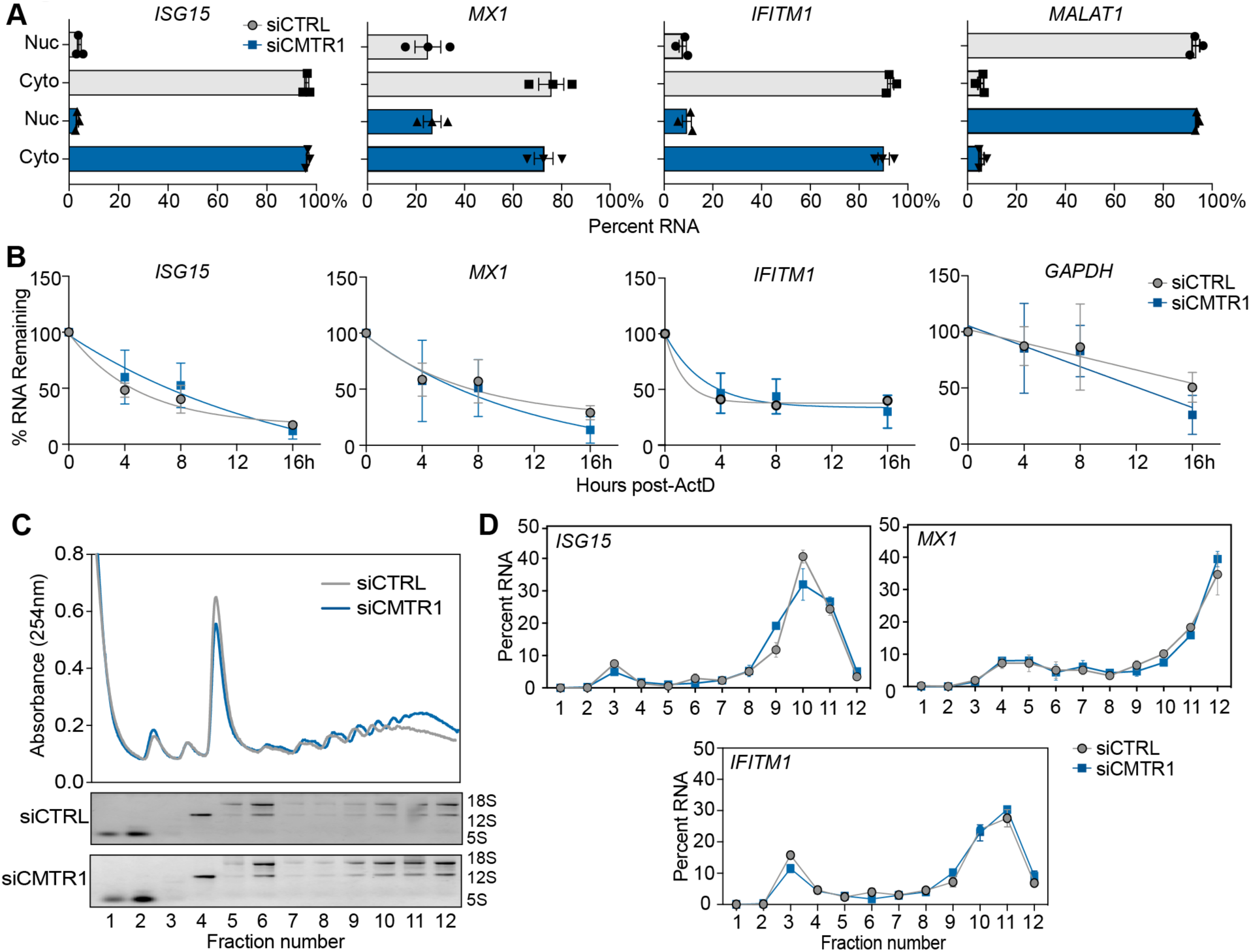
CMTR1 does not regulate the nuclear export, RNA stability, or polysome association of specific ISGs. **(A)** The relative percent of *ISGs* or *MALAT1* (nuclear fractionation control) in nuclear and cytoplasmic fractions of siRNA and IFN-β-treated Huh7 cells, as analyzed by RT-qPCR.**(B)** RNA stability, as measured by the percent of transcript remaining in siRNA and IFN-β-treated Huh7 cells at the indicated times following actinomycin D (10 µg/mL) treatment.**(C)** (Top) A representative plot of the relative absorbance values of fractions isolated from extracts of siRNA and IFN-β-treated Huh7 cells following centrifugation over 15-50% sucrose gradients. (Bottom) RNA from each fraction was separated on an agarose gel and visualized with ethidium bromide. Ribosomal RNA bands (18S, 12S, and 5S) are indicated. **(D)** RT-qPCR analysis of ISGs from polysome profiling of extracts from (C). Data are presented as the percent of total mRNA in each fraction. All IFN-β treatments were performed at 50 U/mL for 6 h. Values are the mean ± SEM of 3 biological replicates.

### CMTR1 inhibits IFIT1-mediated translational regulation of specific ISGs via their 5’ UTRs

mRNAs that lack Cap 1 2’O methylation are known to be preferentially bound by IFIT1 (13, 19-22, 25, 27, 30, 42), and the translation of viral RNAs lacking Cap 1 2’O methylation is known to be inhibited by IFIT1 (20, 21, 42). Therefore, we hypothesized that the protein expression of host transcripts lacking Cap 1 2’O methylation may be inhibited by IFIT1. Indeed, we found that depletion of IFIT1 rescued the protein expression of ISG15, MX1, and IFITM1 resulting in equivalent expression of ISGs following CMTR1 depletion as compared to control in IFN-β-treated Huh7 cells (**Figure 4-B**). We did not examine IFIT3 in this assay, as it is in a complex with IFIT1 alterations in IFIT1 levels likely also IFIT3 expression (20, 21, 43). Therefore, the translational repression of the CMTR1-dependent ISGs is mediated through the actions of IFIT1.

**Figure 4:**
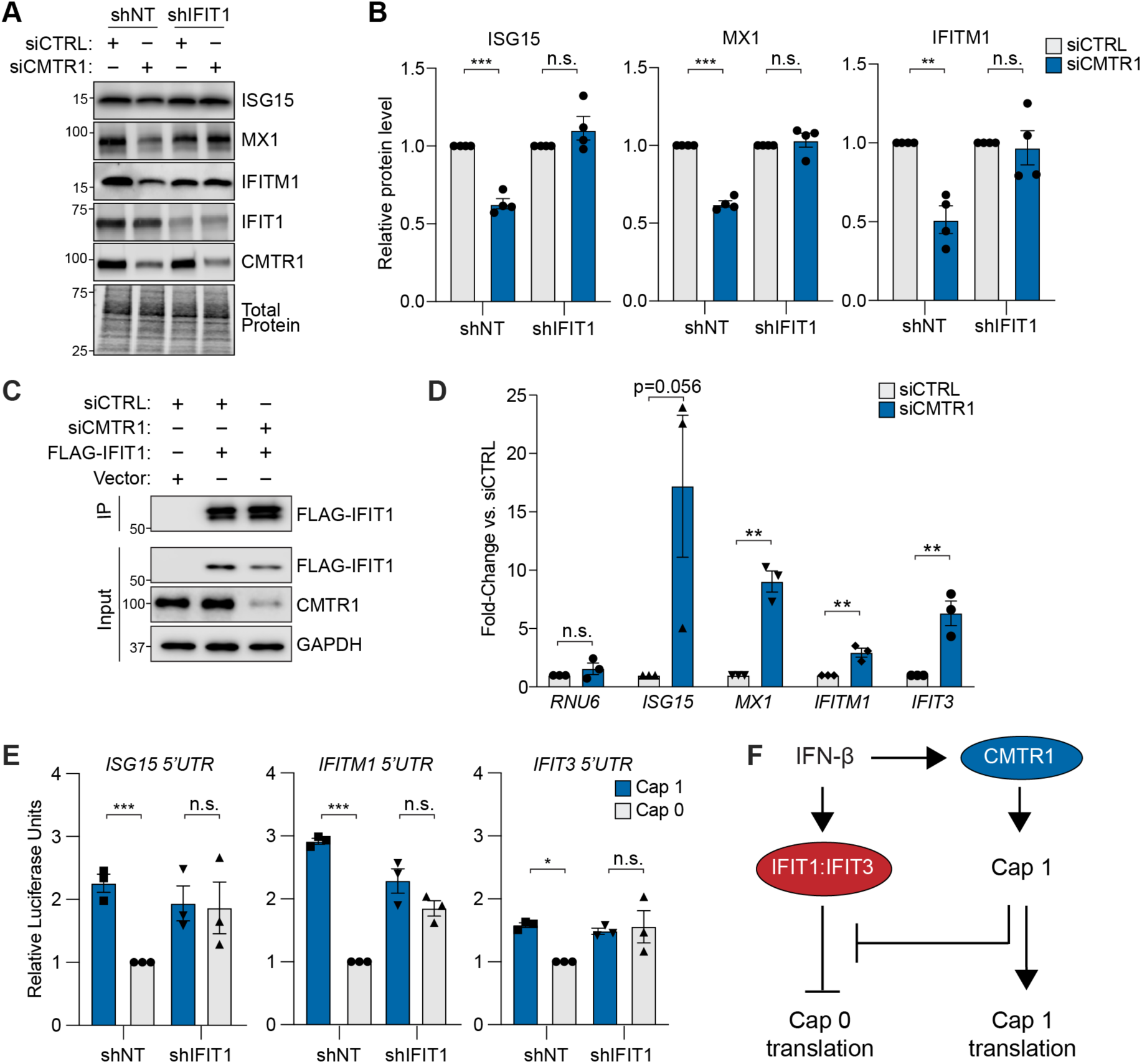
CMTR1 inhibits IFIT1-mediated translational regulation of specific ISGs via their 5’ UTRs. **(A)** Representative immunoblot of extracts of IFN-β-treated Huh7 cells transduced with the indicated shRNA (non-targeting control (NT) and IFIT1) and then transfected with siCTRL or siCMTR1. **(B)** Quantification of immunoblots from (A). The values display the relative protein levels of the indicated ISG normalized to the total protein, which were then set to 1 for each set of siCTRL samples. They represent the mean ± SEM of 4 biological replicates. **(C)** Representative immunoblot of FLAG-immunoprecipitated (IP) or input fractions of IFN-β-treated Huh7 cells transfected with Vector or FLAG-IFIT1, as well as siCTRL or siCMTR1. **(D)** RT-qPCR analysis of enriched transcripts relative to input by anti-FLAG IP in siRNA and IFN-β-treated Huh7 cells. Fold enrichment values for each transcript are graphed relative to siCTRL and are the mean ± SEM of 3 biological replicates. **(E)** Secreted *Gaussia* luciferase units of a reporter whose expression is driven by the 5’ UTR of the indicated ISGs and contains either Cap 0 or Cap 1 in IFN-β-treated Huh7 cells transduced with shNT or shIFIT1. Fold change values for the indicated reporters are shown, with the shNT cells transfected with the indicated Cap 0 construct set to 1 for each experiment, and graphed as the mean ± SEM of 3 biological replicates. (**F**) Following IFN-β-signaling, IFIT1 and IFIT3 are induced, and they can inhibit the translation of Cap 0 mRNAs. However, CMTR1 induction by IFN-β can ensure that the mRNA of ISGs is modified by Cap 1, ensuring ISG protein expression and the antiviral response. For (B) and (D), ** p < 0.01, *** p < 0.005 by unpaired Student’s t test. For (E), * p < 0.05, *** p < 0.005 by ordinary one-way ANOVA with Sidak’s multiple comparison test. n.s. = not significant. All IFN-β treatments were performed at 50 U/mL for 6 h.

To determine how IFIT1 mediates this repression of protein expression following CMTR1 depletion, we first tested if IFIT1 bound selectively to the transcripts of the CMTR1-regulated ISGs. However, we found that IFIT1 bound to both the CMTR1-regulated and -nonregulated transcripts following CMTR1 depletion (**Figure 4C-D**). As a control, we tested IFIT1 binding to the U6 snRNA (*RNU6*), which has a gamma-monomethyl 5’ cap, and we found, similar to others, that IFIT1 did not bind this RNA (44). We next tested if CMTR1-regulated ISGs contained RNA elements that could dictate their sensitivity to translational repression by IFIT1. We generated reporters in which the expression of *Gaussia* luciferase was driven by the 5’ UTRs of either the CMTR1-regulated *ISG15* and *IFITM1*, or the non-CMTR1-regulated *IFIT3*. We used *in vitro* transcription to generate reporter transcripts and then to the first transcribed nucleotide added either the m^7^G modification alone (Cap 0) or added both m^7^G and 2’O methylation of the ribose (Cap 1). These transcripts containing different 5’ UTRs and cap structures were then transfected into IFN-β-primed Huh7 cells. At six hours post transfection, we measured the luciferase activity of these reporters and found that the reporters containing the 5’ UTRs of the CMTR1-regulated ISGs *ISG15* and *IFITM1* had decreased luciferase activity of about 2-3 fold in the absence of 2’O methylation in the cap (Cap 0), while the luciferase activity of the reporter containing the 5’ UTR of the CMTR1-nonregulated ISG *IFIT3* was only modestly altered (decreased 1.5 fold) by lack of Cap 1 2’O methylation (**Figure 4E**). Additionally, we found that when IFIT1 was depleted from these cells, the luciferase activity of these reporters was no longer altered by lack of Cap 1 2’O methylation (**Figure 4E**). Thus, these results reveal that the presence of Cap 1 within the 5’ UTRs of CMTR1-regulated ISGs alleviates the translational inhibition by IFIT1 during the IFN response (**Figure 4F**). Further, these results reveal that the 5’ UTR of the non-CMTR1 regulated ISG IFIT3 contains RNA element(s) that allow it to partially overcome IFIT1-mediated translational inhibition.

## Discussion

We hypothesized that ISGs induced during the IFN response would be reliant on Cap 1 2’O methylation by CMTR1 for evasion of IFIT1 translational inhibition. Here, we show that the protein expression of specific ISGs (ISG15, IFITM1, MX1) is enhanced by CMTR1, while the expression of the ISGs IFIT1 and IFIT3, which suppress translation of both viral and cellular mRNAs lacking Cap 1 2’O methylation (13, 18-22, 25, 27, 30, 45-47), is not as sensitive to CMTR1 levels. Surprisingly, while IFIT1 bound to all tested ISGs when CMTR1 was depleted, only the protein expression of the CMTR1-regulated ISGs was inhibited by IFIT1 in these conditions. Importantly, we found that the 5’ UTR of *IFIT3* was not as reliant as the 5’ UTRs of *ISG15* and *IFITM1* on Cap 1 2’O methylation to prevent IFIT1-mediated translational regulation. Therefore, these data reveal that during the type I IFN response, the mRNAs of some ISGs require CMTR1 to evade sensing and translational inhibition by IFIT1 and that other ISGs encode additional mechanisms to evade this sensing. Ultimately, this suggests that during the IFN response, Cap 1 2’O methylation of transcripts by CMTR1 ensures their protein expression to program a functional IFN response.

We found that CMTR1 is required for the protein expression of ISGs by preventing IFIT1-mediated repression. While others have found that loss of CMTR1 and/or Cap 1 2’O methylation can trigger RIG-I sensing of host mRNAs to induce IFN and ISGs under certain conditions (12, 16), we did not observe this upon CMTR1 depletion in Huh7 cells, which allowed us to focus on how changes in CMTR1 levels affects IFIT1 sensing of RNAs lacking Cap 1 2’O methylation (**Figure 1D**). While the molecular mechanisms by which IFIT1 inhibits the translation of RNAs lacking Cap 1 2’O methylation are not fully clear (13, 18-22, 29, 45-47), it has been shown that IFIT1 can compete with the cap binding protein eIF4E and also with eIF4F for binding to Cap 0 RNAs and prevent formation of the pre-initiation complex (19, 22, 46, 47). Further, it has been shown that murine IFIT1 decreases the translation rate of viral RNAs lacking Cap 1 (19). To date, no study has shown the effects of IFIT1 on the polysome occupancy of its regulated transcripts. Indeed, we did not observe a change in the polysome occupancy of the IFIT1-regulated ISGs upon CMTR1 depletion, even though the reduction in their protein expression upon CMTR1 depletion or by loss of Cap 1 2’O methylation was mediated by IFIT1 **(Figure 4)**. While we do not yet know the mechanism of IFIT1-mediated translational repression, our data support the conclusion that IFIT1 inhibits the translation of CMTR1-regulated ISGs.

Interestingly, the protein expression of IFIT1 and IFIT3 were not significantly affected by CMTR1 depletion, suggesting that these mRNAs encode a mechanism to overcome the restriction that resulted in reduced protein expression of ISG15, MX1, and IFITM1 following CMTR1 depletion. Originally, we hypothesized that the reason IFIT1 and IFIT3 protein expression were not affected by CMTR1 depletion was that these mRNAs were not bound by IFIT1 in the absence of CMTR1. However, we found that *IFIT3* mRNA still bound to IFIT1 in the absence of CMTR1 (**Figure 4**). Therefore, we next tested if 5’ UTR encoded elements of ISGs affected IFIT1-mediated translational inhibition and CMTR1-dependence. Indeed, we found that when IFIT1 was upregulated by IFN treatment, the translation of a luciferase reporter driven by the 5’ UTR of *IFIT3* (not regulated by CMTR1) was only partially inhibited by the absence of Cap 1 2’O methylation, while the translation of reporters with either the 5’ UTR of *ISG15* or *IFITM1* (regulated by CMTR1) was more strongly inhibited by the absence of Cap 1 2’O methylation **(Figure 4)**. This suggests three possible mechanisms that underlie how the CMTR1 sensitivity of these ISGs is regulated. First, it is possible that features in the 5’ UTR of the non-CMTR1 regulated ISG IFIT3 prevent IFIT1 translational inhibition. In support of this idea, the 5’ UTR of some viral RNAs can encode secondary structures that prevent IFIT1 sensing and translational inhibition (18, 30, 42). Alternatively, the non-CMTR1 regulated ISG IFIT3 may only have low levels of Cap 1, and so it may not be as sensitive to changes in CMTR1 and resulting decrease in Cap 1 levels that would enable IFIT1 sensing and translational inhibition. Indeed, CMTR1 has reduced activity on mRNAs with highly structured 5’ UTRs and as such these structured 5’ UTRs require unwinding by the helicase DHX15 helicase for efficient Cap 1 formation (41, 48). Finally, it is possible that CMTR1 depletion preferentially affects the levels of Cap 1 in ISGs with highly structured 5’ UTRs which are not efficiently 2’O methylated, thereby specifically resulting in reduced translation of these transcripts following CMTR1 depletion. In any case, our results suggest that RNA features in ISG transcripts can influence IFIT1 translational regulation. Future work should determine the complete repertoires of CMTR1-dependent and -independent cellular RNAs during the IFN response to define novel elements that mediate the function of IFIT1. Overall, this work reveals that CMTR1 promotes the protein expression of specific ISGs to efficiently establish an IFN-dependent antiviral state.

## Acknowledgements

We thank Michael McFadden and Dr. Christopher Holley for experimental help and advice, colleagues who provided reagents (see Methods), the Duke Functional Genomics Core Facility, and Horner lab members for discussion during preparation of this manuscript. This work was supported by funds from: Burroughs Wellcome Fund (S.M.H.); NIH R01AI125416, R21AI129851 (S.M.H.); NIH F32AI145180 (G.D.W); NIH T32-CA009111 (D.L.S); American Heart Association Pre-doctoral Fellowship, 17PRE33670017 (N.S.G.)

## Author contributions

Conceptualization: N.S.G., G.D.W., D.L.S., S.M.H. Investigation: G.D.W., N.S.G., D.L.S. Formal analysis: G.D.W., N.S.G., S.M.H. Writing: G.D.W., N.S.G., D.L.S., S.M.H. Funding acquisition: G.D.W., N.S.G., D.L.S., and S.M.H.

## Methods

### Cell culture, viral stocks, and viral infection

Huh7, 293T, and Vero cells were grown in Dulbecco’s Modification of Eagle’s Medium (DMEM; Mediatech) supplemented with 10% fetal bovine serum (FBS; HyClone), 25 mM 4-(2-hydroxyethyl)-1-piperazineeethanesulfonci acid (HEPES; Thermo Fisher), and 1X non-essential amino acids (Gibco) further referred to as cDMEM. THP1 cells (gift from Dr. Dennis Ko, Duke University, who obtained them from the American Type Culture Collection (ATCC)) were grown in Roswell Park Memorial Institute medium 1600 (RPMI; Thermo Fisher) supplemented with 10% FBS, and 25 mM HEPES. All cell lines were grown at 37°C supplemented with 5% CO_2_. 293T and Vero (CCL-3216 and CCL-81) cells were obtained from ATCC; Huh7 cells were a gift of Dr. Michael Gale Jr. at the University of Washington; Huh7-STAT1^KO^ have been described previously (49). Generation of shRNA IFIT1 cells: 293T cells were co-transfected with psPax2 and pMD2.G (Addgene plasmids #12260 and #12259), as well as pLKO.1 (50) containing either non-targeting or IFIT1 shRNA (Sigma, TRC1, Clone: TRCN0000158439). Huh7 cells were transduced with the indicated shRNA expressing lentivirus then selected with 2 μg/ml puromycin (Sigma). After selection, cells were maintained in 1 μg/ml puromycin until use. The identity of cell lines was verified using the Promega GenePrint STR kit (DNA Analysis Facility, Duke University), and cells were verified as mycoplasma free by the LookOut Mycoplasma PCR detection kit (Sigma).

Infectious stocks of a cell culture-adapted strain of DENV2-NGC and ZIKV-DAKAR were generated and titered, as described (51). ZIKV-DAKAR (Zika virus/A,africanus-tc/SEN/1984/41525-DAK) was provided by Dr. Scott Weaver at University of Texas Medical Branch. DENV2-NGC and ZIKV-DAKAR infections (MOI 0.01) were performed in Huh7 cells in serum-free media for 4 hours, after which cDMEM was replenished for a total infection time of 48 hours. VSV-GFP, provided by Dr. Sean Whelan at Harvard University, was propagated in Vero cells, as described previously (39). Infections (MOI 0.2) were performed in Huh7 cells in serum-free media for 4 hours, after which cDMEM was replenished for a total infection time of 8 hours. Sendai virus (SenV) Cantell strain was obtained from Charles River Laboratories and used at 200 hemagglutination units/mL. SenV infections were performed in serum-free media for 1 hour, after which cDMEM was replenished.

### Plasmids

The following plasmids used in this study have been previously described: pEF-Tak-Flag (52) and pCMV-Gluc2 (NEB). The following plasmids were generated in this study:

pEF-Tak-Flag-IFIT1, pCMV-5’UTR ISG15 TATT, pCMV-5’UTR IFITM1 TATT, and pCMV-5’UTR IFIT3. To generate pEF-Tak-Flag-IFIT1, HEK293T cells were stimulated with SenV for 24 hours, cDNA was created using Superscript III (Thermo Fisher), and then IFIT1 was amplified by PCR. The IFIT1 amplicon was inserted into the *NotI-PmeI* digested pEF-Tak-Flag vector by InFusion cloning (Clontech). The following 5’UTR constructs were purchased from IDT as gBlocks ISG15 (RefSeq: NM_005101.4; UTRDB: 5HSAR051845), IFITM1 (RefSeq: NM_003641.4; UTRDB: 5HSAR055561), and IFIT3 (RefSeq: NM_001549.6; UTRDB: 5HSAR021528) and inserted into *HindIII-BamHI* digested pCMV-Gluc2 by InFusion cloning ((53); http://utrdb.ba.itb.cnr.it). This resulted in pCMV-5’UTR IFIT3. For ISG15 and IFITM1, the final nucleotide of the TATA box of pCMV-Gluc2 was changed to a T by using site-directed mutagenesis (QuikChange lightning; Agilent) to facilitate *in vitro* transcription (54), resulting in pCMV-5’UTR ISG15 TATT and pCMV-5’UTR IFITM1 TATT. All DNA sequences were verified by sequencing. Primer sequences are provided in **Table 1**.

**Table 1:**
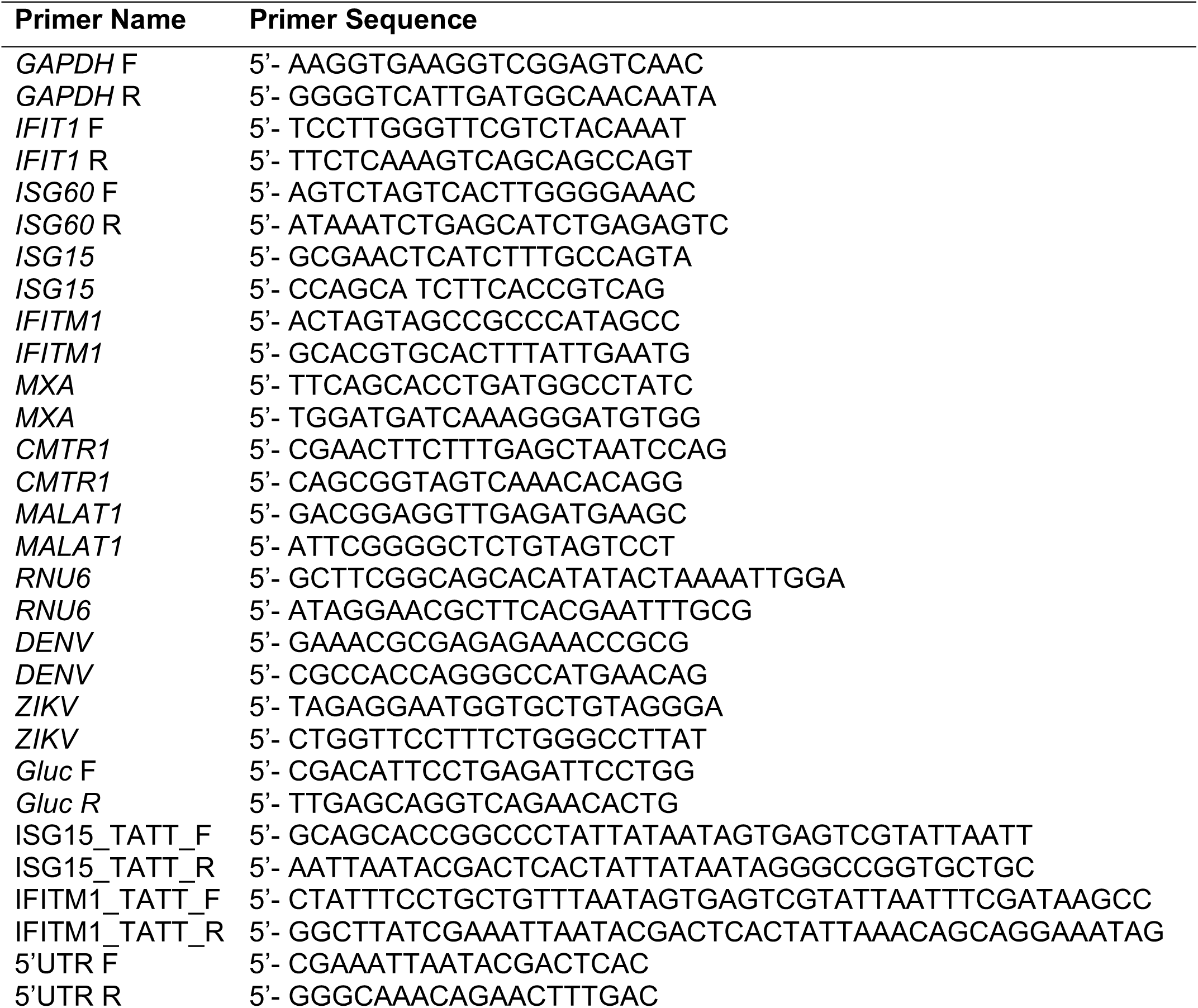
Primers used for cloning and RT-qPCR.

### *In vitro* transcription and mRNA capping

Purified *NotI*-linearized pCMV-Gluc2 DNA was used as a template for *in vitro* transcription using the MEGAscript T7 transcription kit (Ambion). The *in vitro* transcribed RNA was treated with DNase I (Ambion) and then purified by phenol-chloroform-isoamyl acid (Thermo Fisher) and subsequent isopropanol precipitation. Purified RNA was treated with the Vaccinia Capping Enzyme (NEB) to generate Cap 0 and in some cases also the Vaccinia 2’ O methyltransferase (NEB) to generate Cap 1. The quality of the RNA was verified by denaturing agarose gel electrophoresis.

### Transfection

DNA transfections were performed using PEI MAX (Polysciences, Inc.). *In vitro* transcribed RNAs were transfected using TransIT-mRNA transfection kit (Mirus). siRNAs directed against CMTR1 (Dharmacon, Horizon Discovery, M-014142-00-0005) or non-targeting siRNA Pool #1 (Dharmacon, Horizon Discovery, D-001206-13-05) were transfected using Lipofectamine RNAiMax (Invitrogen), and experiments were performed at 48 hours post-transfection.

### IFN-β treatment

All IFN-β (PBL Assay Science) treatments were performed at a concentration of 50 U/mL in cDMEM, unless otherwise noted.

### Luciferase assay

*Gaussia* luciferase activity was measured in supernatants harvested from Huh7-shCTRL or Huh7-shIFIT1 cells that had been transfected with *in vitro* transcribed RNA (6 hours) after IFN-β treatment (6 hours), according to the manufacturer’s instructions (Gaussia Luciferase Flash Assay Kit; Pierce). Samples were read on the BioTek Synergy 2 multi-mode microplate reader. Data represent the average of two technical replicates performed for each biological replicate.

### Focus-forming assay

Serial dilutions of supernatants were used to infect naïve Vero cells in triplicate wells of a 48-well plate. Infected cells were then overlaid with methylcellulose. At 48 hours post-infection, cells were fixed in cold 1:1 methanol:acetone and immunostained with anti-flavivirus 4G2 antibody that recognizes the E protein that was purified in the lab from a hybridoma (1:2000). Following binding of horseradish peroxidase-conjugated secondary antibody (1:1000; Jackson ImmunoResearch), infected foci were visualized with the VIP Peroxidase Substrate Kit (Vector Laboratories) and counted at 40X magnification. Titer was calculated using the following formula: (dilution factor x number of foci x 1000) / volume of infection (μl), resulting in units of focus forming units / mL.

### Quantification of viral infection by fluorescence microscopy

For DENV and ZIKV infections, Huh7 cells treated with siRNAs were infected with DENV or ZIKV (MOI 0.01). At 48 hours post-infection, these cells were fixed in 4% paraformaldehyde in PBS, permeabilized with 0.2% Triton X-100 in PBS, blocked with 3% BSA in PBS, and then immunostained with anti-flavivirus 4G2 antibody (1:1000). Infected cells were visualized by AlexaFluor 488-conjugated secondary antibody (1:1000, Thermo Fisher) and nuclei stained with DAPI (Thermo Fisher). For VSV infections, Huh7 or STAT1^KO^ cells treated with siRNAs were pre-treated with IFN-β (25 U/mL) for 16 hours and then infected with VSV-GFP (MOI 0.2). At 8 hours post-infection, cells were fixed in 4% paraformaldehyde in PBS and nuclei stained with DAPI. Cells were imaged with the Cellomics Arrayscan VTI microscope at the Duke Functional Genomics Core Facility. The percentage of infected cells was determined by measuring cells that stained positive for viral antigen relative to the total number of nuclei (10 fields per well, >5000 cells per condition).

### RT-qPCR

Total cellular RNA was extracted as described in the associated method below or by using the RNeasy RNA mini kit (Qiagen). Then, the RNA was reverse transcribed using the iScript cDNA synthesis kit (Bio-Rad), as per the manufacturer’s instructions. The resulting cDNA was diluted 1:5 in water. RT-qPCR was performed in triplicate using the Power SYBR Green PCR master mix (Thermo Fisher) and the QuantStudio 6 Flex RT-PCR systems. Primer sequences are listed in **Table 1**.

### Immunoblotting

Cells were lysed in a modified radioimmunoprecipitation assay (RIPA) buffer (50 mM Tris (pH 7.5), 150 mM NaCl, 5 mM EDTA, 0.1% sodium deoxysulfate, 0.5% sodium deoxycholate, and 1% Triton X-100) supplemented with protease inhibitor cocktail (Sigma) and Halt Phosphatase Inhibitor (Thermo Fisher), and post-nuclear lysates were isolated by centrifugation. Quantified protein (7-10 µg) was resolved by SDS/PAGE, transferred to nitrocellulose membranes in the following buffer: 25 mM Tris-HCl, 192 mM glycine, 0.01% SDS, then blocked in 3% Bovine Serum Albumin (Sigma, BSA) in Tris-buffered saline containing 0.01% Tween-20 (TBS-T). After washing with phosphate buffered saline containing 0.01% Tween-20 (PBS-T) or TBS-T (for p-STAT1) buffer, membranes were incubated with species-specific horseradish peroxidase-conjugated antibodies (Jackson ImmunoResearch, 1:5000) followed by treatment of the membrane with Clarity Western ECL substrate (Bio-Rad) and imaging on a LICOR Odyssey FC. The following antibodies were used for immunoblot analysis: rabbit anti-CMTR1 (Atlas Antibodies, HPA029980, 1:1000); mouse anti-ISG15 (Santa Cruz Biotechnology, sc-166755, 1:1000); mouse anti-IFITM1 (Proteintech, 60074-Ig, 1:1000); mouse anti-Tubulin (Sigma, T5168, 1:5000); rabbit anti-MX1 (Abcam, ab207414, 1:1000); mouse anti-IFIT3 (Abcam, ab76818, 1:1000); rabbit anti-IFIT1 (gift of Dr. Ganes Sen, (17)); mouse anti-pSTAT1 (pY701; BD, 612132, 1:1000); rabbit anti-GAPDH (GeneTex, GT239, 1:1000) mouse anti-STAT1 (BD, 610115, 1:1000); mouse anti-FLAG (Sigma, F3165, 1:5000).

### Quantification of immunoblots

Following imaging using the LI-COR Odyssey Fc, immunoblots were quantified using ImageStudio Lite software, and raw values were normalized to total protein (Revert 700 total protein stain; LI-COR) for each condition.

### Polysome profiling

Cells were harvested from 10 cm plates by trypsinization at indicated time points (0.2 mM; Sigma) and were lysed in the following lysis buffer (200 mM KCl, 25 mM HEPES (pH 7.0), 10 mM MgCl_2_, 2% n-Dodecyl β-D-maltoside (Chem-Impex), 1 mM DTT, 40 U RNaseIn) for 5 minutes on ice. Post-nuclear lysates were isolated and then centrifuged on 15-50% sucrose gradients prepared in polysome gradient buffer (200 mM KCl, 25 mM HEPES (pH 7.0), 15 mM MgCl_2_, 1 mM DTT) at 35,000 xG for 3.5 hours at 4°C. Following centrifugation, 12 fractions were collected from each sample using a Bio Comp piston gradient fractionator fitted with a TRIAX flow cell to measure absorbance. RNA was extracted from each fraction using TRIzol LS (Thermo Fisher), and RNA quality was assessed on a 1% agarose gel. Equal volumes of RNA from each fraction were subjected to RT-qPCR.

### Immunoprecipitation of IFIT1-bound RNA

Cell extracts were harvested in lysis buffer (100 mM KCl, 5 mM MgCl_2_, 10 mM HEPES (pH 8.0), 0.5% NP-40 supplemented with protease inhibitor cocktail (Sigma) and RNasin ribonuclease inhibitor (Promega)), and lysates were cleared by centrifugation. RNP complexes were immunoprecipitated with Protein G Dynabeads (Invitrogen) beads previously bound with anti-FLAG antibody (Sigma) overnight at 4°C with head-over-tail rotation, and then washed five times in ice-cold NT2 buffer (50 mM Tris-HCl (pH 7.4), 150 mM NaCl, 1 mM MgCl_2_, 0.05% NP-40). Protein for immunoblotting was eluted from 25% of beads by heating at 95°C for 5 minutes in 2X Laemmli sample buffer (Bio-Rad). RNA was extracted from the remaining 75% of beads using TRIzol (Thermo Fisher). Equal volumes of eluted RNA were used for cDNA synthesis, quantified by RT-qPCR, and normalized to RNA levels of input samples.

### Nuclear-cytoplasmic fractionation

Following siRNA treatment (48 hours) and IFN-β treatment (6 hours), cells were harvested by trypsinization and lysed in 200 µL lysis buffer (10 mM Tris-HCl (pH 7.4), 140 mM NaCl, 1.5 mM MgCl_2_, 10 mM EDTA, 0.5% NP-40) on ice for 5 minutes. Following centrifugation at 12000 xG at 4°C for 5 minutes, the supernatant (cytoplasmic fraction) was collected, and the nuclear pellet was rinsed twice with lysis buffer. RNA was extracted from cytoplasmic and nuclear fractions using TRIzol and analyzed by RT-qPCR.

### Measurement of mRNA stability

Following siRNA treatment (48 hours) and IFN-β treatment (6 hours), cells were treated with 10 µg/mL actinomycin D (Sigma). Lysates were harvested at the indicated time points post-treatment and RNA was extracted using TRIzol and analyzed by RT-qPCR. Data were normalized as the percent of RNA remaining at each time point after treatment, relative to that at the time of treatment.

### Statistical Analysis

Statistical tests were performed using GraphPad Prism Version 8.3.0.

